# Calbindin Stratifies Midbrain Dopaminergic Neurons Governing Distinct Aspects of Locomotion

**DOI:** 10.1101/2025.07.07.663227

**Authors:** Cyril Bolduc, Cameron Oram, Skylar Donovan, Haleigh Bach, Martha Liu, Rafaëlle Marier, Morgan Sharpe, Cédric Campeau, Carl Duncan Spencer, Sarah A. Martin, Rajeshwar Awatramani, Jean-François Poulin

## Abstract

Despite advances in delineating the molecular diversity and projection patterns of midbrain dopaminergic (DA) neurons, their specific contributions to locomotion and motor learning remain poorly defined. Here, we applied intersectional ablation and chemogenetic approaches to dissect the distinct roles of calbindin-expressing (CALB1^+^) and non-expressing (CALB1^-^) DA neurons in locomotion. Using newly engineered intersectional constructs, we ablated CALB1^+^ or CALB1^-^ DA neurons in the mouse midbrain. Loss of either subtype led to pronounced deficits in the initiation and vigor of voluntary movements, as demonstrated by a reduction in peak speed, acceleration and deceleration of locomotor bouts. Notably, only CALB1^-^ ablation disrupted locomotor learning. Beyond these functional effects, we observed that selective ablation of CALB1⁺ DA neurons induced local microglial activation and was followed by a non-cell-autonomous loss of CALB1^-^ DA neurons, suggesting that CALB1^-^ neurons are more vulnerable to inflammation triggered by CALB1⁺ neuron loss. We then confirmed these findings by performing acute inhibition of either population using inhibitory DREADD hM4Di. CALB1^-^ DA neurons inhibition impaired the initial acquisition of locomotor learning, whereas inhibition of CALB1^+^ DA neurons disrupted the retention of acquired motor skills from previous days. Moreover, inhibition of CALB1^+^ DA neurons further impaired the initiation and amplitude of voluntary movements, as well as the velocity and acceleration/deceleration of locomotor bouts. Together, these findings provide causal evidence for functional specialization among molecularly distinct midbrain DA subtypes and reveal new aspects of mesostriatal circuit organization underlying locomotion and motor memory.

## INTRODUCTION

In mammals, midbrain dopaminergic (DA) neurons are mainly found within three regions: the substantia nigra pars compacta (SNc), the ventral tegmental area (VTA) and the retrorubral area (RR). These structures play essential roles in movement, motivation, and learning^1–4^. However, previous profiling of sparse-labelled neurons has revealed heterogeneity in efferent projections^5–13^, which suggests that anatomical location alone does not account for their functional diversity. Single-cell RNA sequencing studies across species, including rodents, primates, and humans, have highlighted the molecular and spatial diversity of midbrain DA neurons^14–27^. Additionally, genetic approaches have demonstrated that molecularly defined DA subtypes have distinct projection patterns^25,28–38^, indicating that they may be integrated into specialized functional circuits. Despite these advances, the functional contributions of individual DA subtypes remain poorly understood.

Locomotion, which refers to the ability of an animal to propel its body from one location to another using movements, critically depends on DA signaling. Degeneration of DA neurons underlies bradykinesia in Parkinson’s disease (PD)^39–41^. Notably, not all DA neurons are equally vulnerable to degeneration. Studies show selective patterns of DA axon loss in the striatum, and a more pronounced neuron loss in the SNc compared to the VTA^41–47^. Within the SNc, degeneration is most severe in the ventrolateral tier, which is enriched in ALDH1A1, SOX6, and ANXA1 DA neurons^33,48–53^. Conversely, DA neurons expressing CALB1 are notably more resilient to degeneration, particularly in the VTA and dorsal tier of the SNc^21,33,53–58^. However, it remains unclear whether molecularly-defined DA neuron populations serve distinct roles in locomotion.

Recent studies have begun to elucidate the functional roles of various DA subtypes in locomotion. Ablation of ALDH1A1 DA neurons and silencing of ANXA1 DA neurons have been shown to impair the vigor of movements and locomotor learning^28,29,35^. Calcium recordings from ANXA1 DA neurons have revealed a correlation between their activity and acceleration during locomotion, while CALB1 and VGLUT2 DA neurons are more closely associated with deceleration^25^, suggesting that distinct DA subtypes contribute differentially to locomotion. However, causal studies are required to determine if distinct molecularly-defined DA subtypes encode different roles in locomotion. Furthermore, although CALB1 expression delineates DA neuron populations that differ in projection patterns^25,34^, DA release dynamics^59–62^, and vulnerability to degeneration in PD^21,33,53–58^, the specific behavioral contributions of CALB1^+^ versus CALB1^-^ DA neurons remain poorly understood. Thus, the functional contribution of these populations on locomotion remains to be addressed.

To address this gap, we engineered novel intersectional tools that enable selective ablation of CALB1^-^ and CALB1^+^ DA neurons using intronic recombinase sites enabling combinatorial targeting (INTRSECT)^63,64^. We found that ablation of either population impaired locomotor initiation and vigor, as evidenced by reduced peak speed, acceleration, and deceleration during locomotor events. Notably, ablation of CALB1^-^ DA neurons, but not CALB1^+^ DA neurons, resulted in a locomotor learning deficit. To assess whether permanent ablation can lead to compensatory changes obscuring the contributions of specific DA subtypes to locomotion, we next employed a chemogenetic approach to acutely inhibit individual populations. We found that inhibiting CALB1^-^ DA neurons impaired initial acquisition of locomotor skills, whereas inhibition of CALB1^+^ DA neurons impaired their retention. Finally, the acute inhibition of either population impaired the locomotor initiation and vigor of movements, but was further impaired when CALB1^+^ DA neurons were inhibited. Together, these findings provide causal evidence that molecularly defined DA subtypes play specialized roles in locomotion, shedding light on mesostriatal circuitry and its potential implications for movement disorders like Parkinson’s disease.

## RESULTS

### Intersectional autocleavable Caspase3 constructs selectively ablates molecularly-defined DA subtypes

Genetic ablation of CALB1^+^ and CALB1^-^ neurons can provide a binary response elucidating their contribution to specific locomotor features. A Tobacco Etch Virus (TEV) protease Autocleavable Caspase3 (taCasp3) has been designed to ablate neuronal populations such as ALDH1A1^+^ DA neurons^35,65^. However, to the best of our knowledge, taCasp3 has not been used to ablate neurons using intersectional genetics.

To ablate molecularly-defined DA subtypes, we used the INTRSECT strategy to generate Cre_OFF_Flp_ON_-taCasp3 and Cre_ON_Flp_ON_-taCasp3 constructs (Figure1A). We encoded the constructs into AAV2/9 viral vectors, injected these into the SNc of Calb1-ires-Cre/Dat-2A-Flp mice along with AAVs encoding Cre_ON_Flp_ON_-EYFP and Cre_OFF_Flp_ON_-mCherry reporters, and assessed the efficiency and specificity of our intersectional constructs (Figure1B). Four weeks post-injection, the brains were collected for histological analyses.

**Figure 1.**
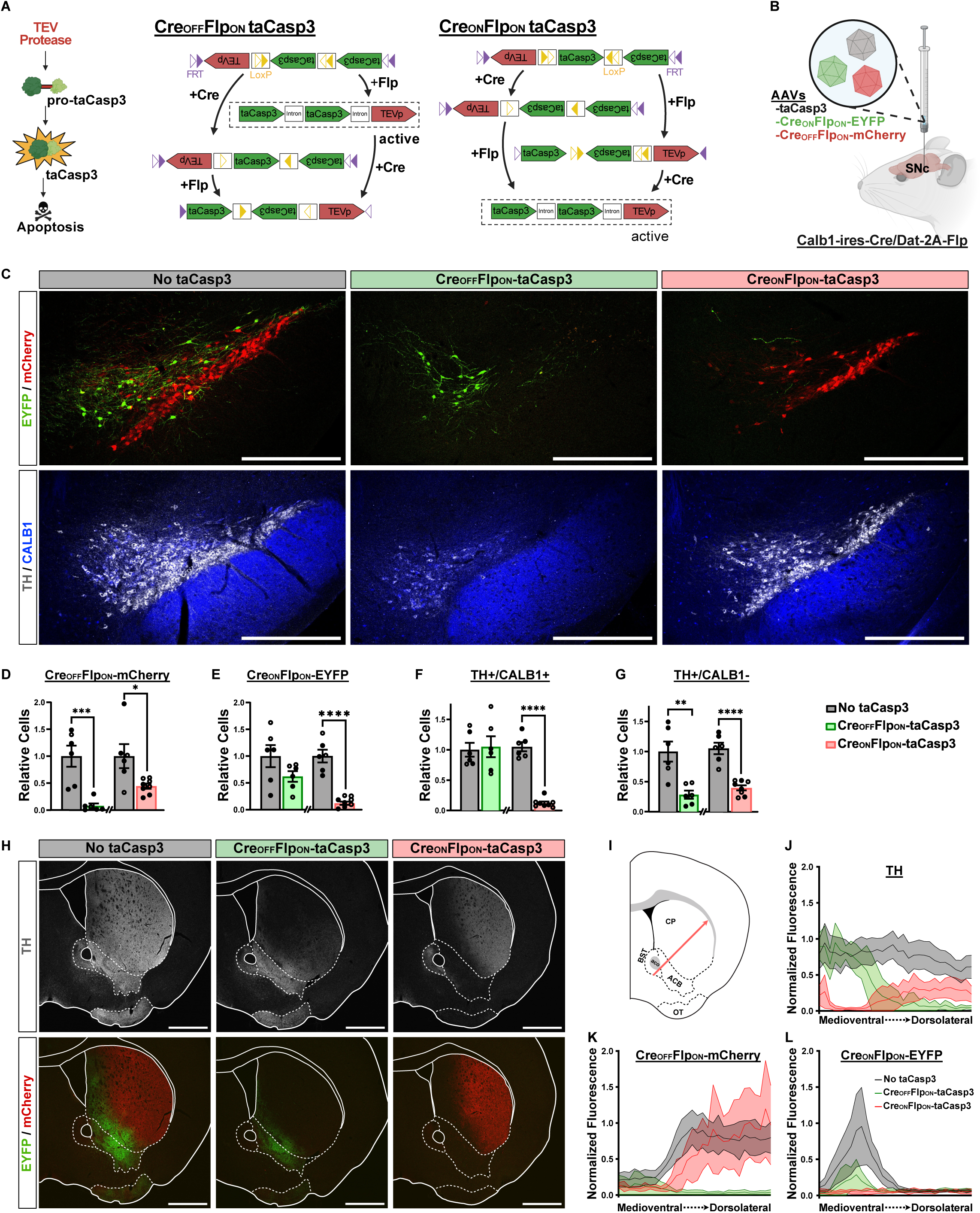
Generation of intersectional autocleavable Caspase3 constructs to selectively ablate targeted molecularly-defined DA subtypes. (A) Cartoon of taCasp3 mechanism of action (adapted from Yang et al.^65^) and schematic of autocleavable Caspase3 INTRSECT design for intersectional ablation.(B) Experimental strategy for the validation of autocleavable Caspase3. Calb1-ires-Cre/Dat-2A-Flp mice were injected in the SNc with or without an AAV encoding taCasp3. Other AAVs encoding Cre_ON_Flp_ON_-EYFP and Cre_OFF_Flp_ON_-mCherry were co-injected to label CALB1^+^ and CALB1^-^ DA neurons respectively. (C) Representative images of midbrain sections co-stained for EYFP/mCherry and TH/CALB1. Scale bar = 500µm. (D) Quantification of Cre_OFF_Flp_ON_-mCherry^+^, (E) Cre_ON_Flp_ON_-EYFP^+^, (F) TH^+^/CALB1^+^, and (G) TH^+^/CALB1^-^ cells in the midbrain relative to the control average. (H) Representative images of striatal sections stained for TH and EYFP/mCherry. Scale bar = 1000µm. (I) Cartoon showing the position of the arrow to quantify the level of denervation from the medioventral to dorsolateral segment of the striatum. (J) Quantification of the medioventral to dorsolateral striatal of the fluorescent level of TH, (K) Cre_OFF_Flp_ON_-mCherry, and (L) Cre_ON_Flp_ON_-EYFP relative to the control average maximal fluorescent intensity value. Filled dots represent the males and empty dots represent the females. Error bars represent the SEM for D-G and 95% Confidence Interval for J-L. Pvalues were determined by a Student T-Test for D-G. *P<0.05, **P<0.01, ***P<0.001, and ****P<0.0001. Abbreviations: ACB = Nucleus accumbens, aco = anterior commissure, BST = Bed nuclei of the stria terminalis, CP = Caudoputamen, OT = Olfactory tubercle, SNc = Substantia nigra pars compacta.

In the absence of taCasp3, we observed a clear segregation of CALB1^+^ and CALB1^-^ DA neurons, consistent with prior studies^25,34^. For instance, CALB1^+^ DA neurons (EYFP^+^), were primarly located to the VTA, dorsal and pars lateralis segments of the SNc, and medial RR. CALB1^-^ DA neurons (mCherry^+^) were concentrated in the SNc, lateral RR, and to a lesser extent in the VTA (FigureS1A-B). Quantification revealed that injection of Cre_OFF_Flp_ON_-taCasp3 resulted in the ablation of 92% of mCherry^+^ and 72% of TH^+^/CALB1^-^cells, while the number of EYFP^+^ and TH^+^/CALB1^+^ cells was not significantly reduced (Figure1C-G). Conversely, injection of Cre_ON_Flp_ON_-taCasp3 ablated 88% of EYFP^+^ and 90% TH^+^/CALB1^+^ cells, but also observed a loss of 59% of mCherry^+^ and 63% of TH^+^/CALB1^-^ cells (Figure1C-G). In both cases, ablation was observed along the full rostrocaudal axis of the midbrain (Figure S1A). To assess the impact on axonal projections, we then looked at the striatal projections of CALB1^+^ and CALB1^-^ DA neurons by quantifying the fluorescence of mCherry^+^ and EYFP^+^ afferents. We found that CALB1^+^ and CALB1^-^ DA neurons differentially innervate the ventromedial and dorsolateral segments of the striatum, as reported previously^25,34^. However, the injection of Cre_OFF_Flp_ON_-taCasp3 resulted in a net decrease in TH^+^ fibers in the dorsolateral segment of the striatum, receiving afferents from the CALB1^-^ DA neurons. In contrast, even though the injection of Cre_ON_Flp_ON_-taCasp3 denervated part of the CALB1^-^ projections to the dorsolateral striatum, it mostly denervated the ventromedial segment receiving inputs from CALB1^+^ DA neurons (Figure1H-L and FigureS1C-L). Together, these results indicate that our intersectional taCasp3 constructs preferentially ablate the targeted molecularly-defined neuronal population and their corresponding striatal projections.

### The absence of off-target ablation and increased microglia density indicates that the nonspecific loss of CALB1^-^ DA neurons is non-cell autonomous

Since we noticed that the injection of our Cre_ON_Flp_ON_-taCasp3 inadvertently killed half of the CALB1^-^ DA neurons, we tested if our constructs had off-target activity. In Calb1-ires-Cre/Dat-2A-Flp mice, the CALB1^-^ DA neurons correspond to the DA neurons that express Flp but not Cre and thus could indicate putative off-target activity in the Flp only configuration. To test this possibility, we co-injected Dat-2A-Flp mice with an AAV encoding a fDIO-mCherry reporter along with AAVs encoding either the Cre_OFF_Flp_ON_ or Cre_ON_Flp_ON_-taCasp3 (Figure2A). Histological analysis revealed that the mCherry^+^ and TH^+^ midbrain neurons and their projections to the striatum were almost completely abolished (99% reduction for mCherry^+^ and 87% for TH^+^) by the injection of Cre_OFF_Flp_ON_-taCasp3 but were left unchanged by the injection of Cre_ON_Flp_ON_-taCasp3 (Figure2B-G). In addition, we also tested the possibility of off-target activity induced by Cre recombination by co-injecting Dat-ires-Cre mice with a DIO-EYFP reporter along with another AAV encoding either the Cre_OFF_Flp_ON_ or Cre_ON_Flp_ON_-taCasp3 constructs (Figure2H). We found that neither the injection of AAV encoding the Cre_OFF_Flp_ON_ or Cre_ON_Flp_ON_-taCasp3 had an impact on the amount of EYFP^+^ and TH^+^ midbrain neurons and their projections to the striatum (Figure2I-N). Together, these results demonstrate the specificity of our intersectional taCasp3 constructs, indicating that the observed loss of CALB1^-^ DA neurons following CALB1^+^ DA neurons ablation is most likely non-cell autonomous.

**Figure 2.**
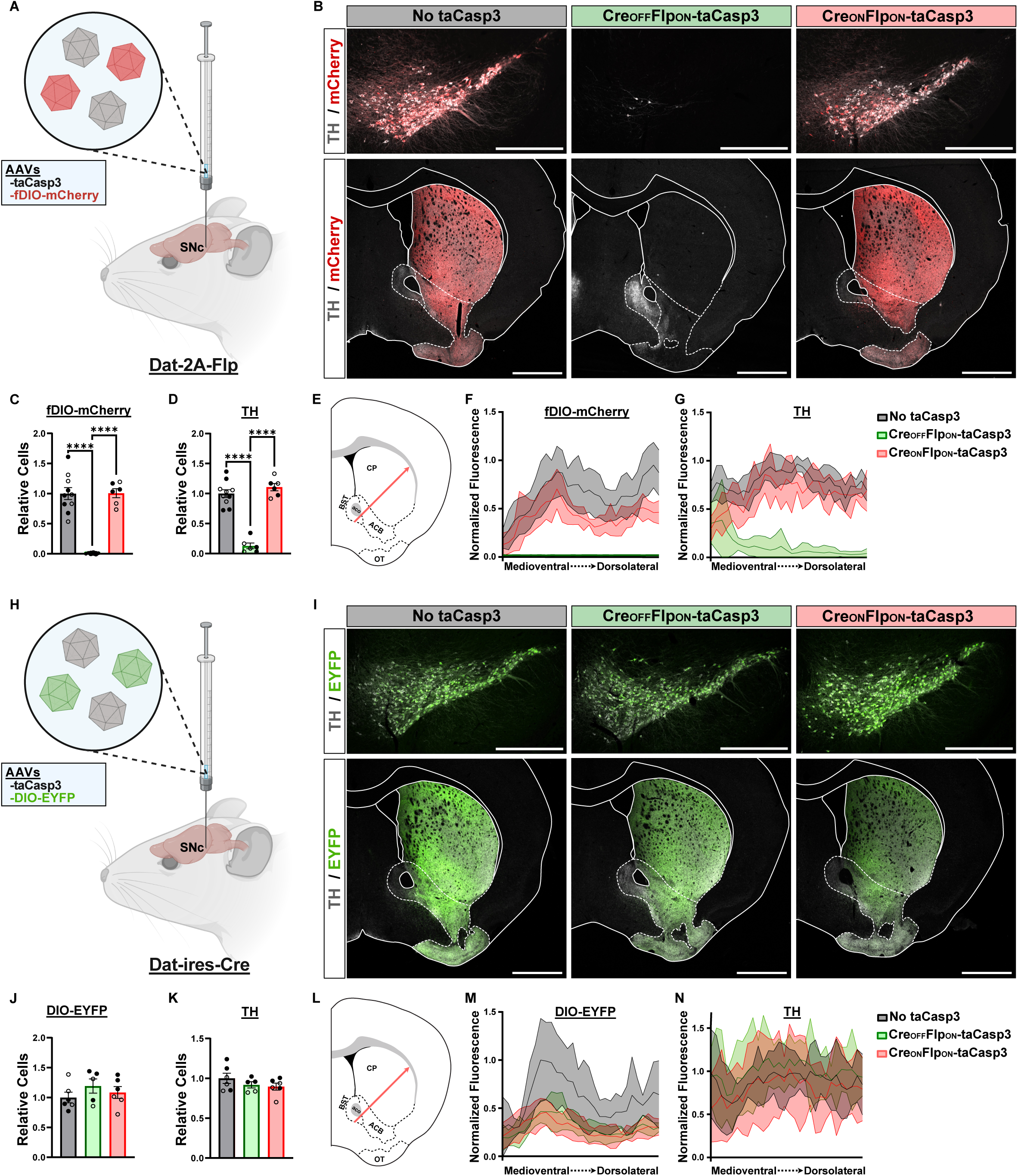
The absence of off-target ablation indicates that the unspecific loss of CALB1^-^ DA neurons is non-cell autonomous. (A) Experimental design for testing off-target activity in Flp expressing neurons only. Dat-2A-Flp mice were injected in the SNc with or without an AAV encoding taCasp3. Another AAV encoding a fDIO-mCherry construct was co-injected to label the infected neurons. Representative images of midbrain and striatal sections co-stained for TH and fDIO-mCherry. Scale bar = 500µm for midbrain images. Scale bar = 1000µm for striatum images. (C) Quantification of fDIO-mCherry^+^ and (D) TH^+^ cells in the midbrain relative to the control average. (E) Cartoon showing the position of the arrow to quantify the level of denervation from the medioventral to dorsolateral segment of the striatum. (F) Quantification of the medioventral to dorsolateral striatal of fDIO-mCherry, and (G) TH fluorescent level relative to the control average maximal fluorescent intensity value. (H) Experimental design for testing off-target activity in Cre expressing neurons only. Dat-ires-Cre mice were injected in the SNc with or without an AAV encoding taCasp3. Another AAV encoding a DIO-EYFP construct was co-injected to label the infected neurons. (I) Representative images of midbrain and striatal sections co-stained for TH and DIO-EYFP. Scale bar = 500µm for midbrain images. Scale bar = 1000µm for striatum images. (J) Quantification of DIO-EYFP^+^, and (K) TH^+^ cells in the midbrain relative to the control average. (L) Cartoon showing the position of the arrow to quantify the level of denervation from the medioventral to dorsolateral segment of the striatum. (M) Quantification of the medioventral to dorsolateral striatal of DIO-EYFP, and (N) TH fluorescent level relative to the control average maximal fluorescent intensity value. Filled dots represent the males and empty dots represent the females. Error bars represent the SEM for C-D and J-K and 95% Confidence Interval for F-G and M-N. Pvalues were determined by a one way ANOVA with Tukey’s post-hoc test for C-D and J-K. *P<0.05, **P<0.01, ***P<0.001, and ****P<0.0001. Abbreviations: ACB = Nucleus accumbens, aco = anterior commissure, BST = Bed nuclei of the stria terminalis, CP = Caudoputamen, OT = Olfactory tubercle, SNc = Substantia nigra pars compacta.

We wondered if inflammation could explain the unexpected loss of CALB1^-^ neurons after ablation of the CALB1^+^ DA neuron population. To explore this possibility, we quantified the density of IBA1^+^ microglia in both the SNc and VTA (FigureS2A). We found that, upon the ablation of either CALB1^-^ or CALB1^+^ DA neurons, the density of IBA1^+^ microglia was increased in the SNc, with a similar trend observed in the VTA, although not significant for Cre_ON_Flp_ON_-taCasp3 (FigureS2B-C). These findings indicate that the ablation of CALB1^+^ DA neurons induces a local inflammation, which might eventually kill the CALB1^-^ DA neurons. It also suggests that CALB1^+^ DA neurons are more resilient to local inflammation than the CALB1^-^ DA neurons.

### Ablation of CALB1^-^ or CALB1^+^ DA neurons differently impairs locomotor learning, initiation, and vigor of ongoing movements

To assess the role of CALB1^-^ and CALB1^+^ DA neurons on locomotion, we first performed a bilateral ablation of each of these populations (Figure3A). The ablations covered the SNc in its entirety and the majority of VTA and RR (FigureS1B). Unfortunately, due to severe weight loss, we had to euthanize 62% of the mice with bilateral ablation of CALB1^-^ DA neurons as early as 5 days post-surgery. This phenotype reflects what is observed after a complete chemical lesion of the SNc^66–70^, but contrasts with the milder phenotype resulting from the ablation ALDH1A1^+^ DA neurons of the SNc^35^. The survival rate increased to 92% if CALB1^-^ DA neurons were ablated unilaterally (FigureS3B-D). On another hand, bilateral ablation of CALB1^+^ DA neurons did not affect the survival rate or body weight (Figure3B-D). These results indicate that CALB1^-^ DA neurons, but not CALB1^+^, are essential for survival.

**Figure 3.**
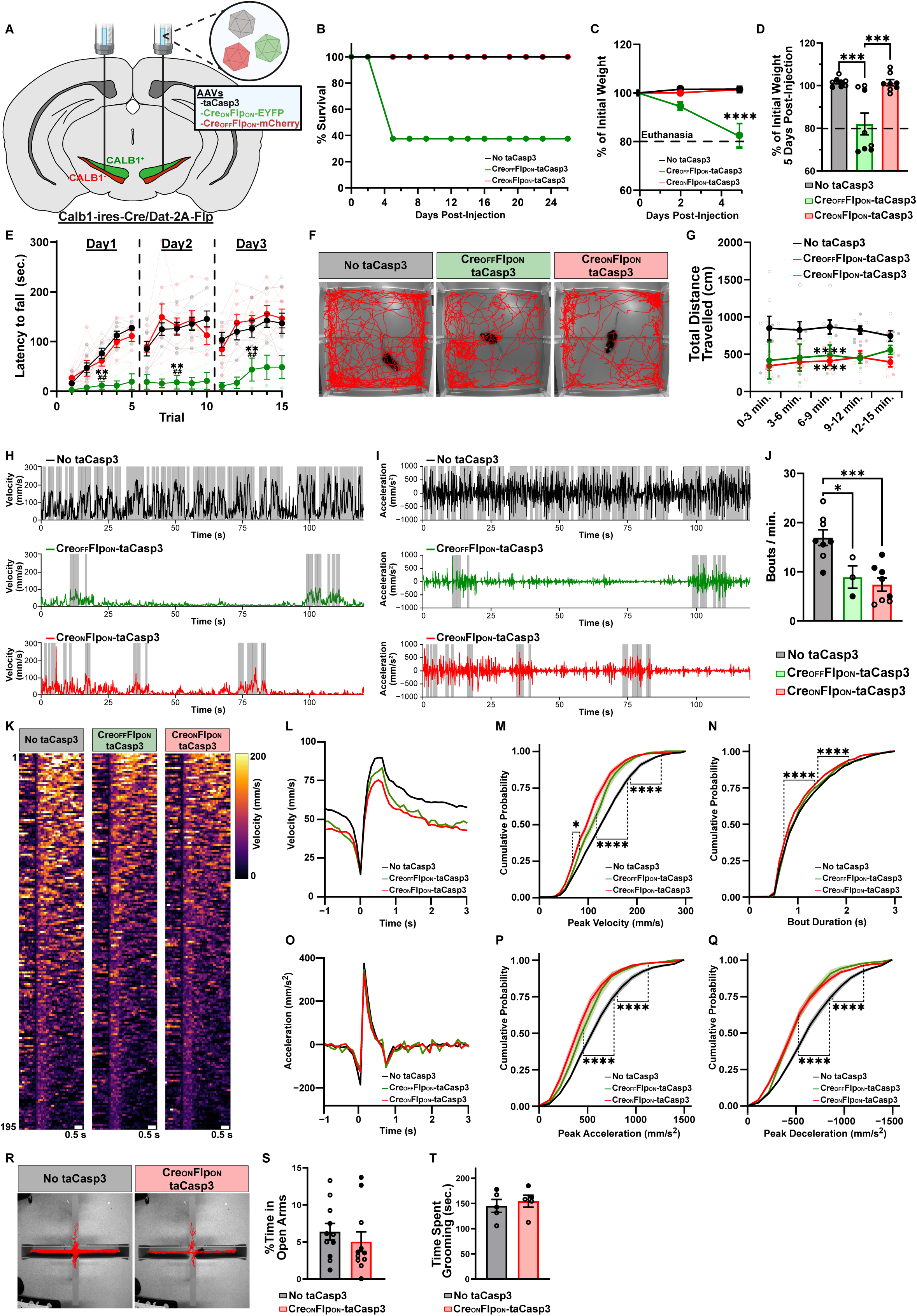
Bilateral ablation of CALB1^-^ or CALB1^+^ DA neurons differently impairs locomotor learning, initiation, and vigor of ongoing movements. (A) Experimental strategy to bilaterally ablate either CALB1^-^ or CALB1^+^ DA neurons. Calb1-ires-Cre/Dat-2A-Flp mice were bilaterally injected of an AAV encoding either Cre_OFF_Flp_ON_ or Cre_ON_Flp_ON_-taCasp3. Other AAVs encoding encoding Cre_ON_Flp_ON_-EYFP and a Cre_OFF_Flp_ON_-mCherry were co-injected. (B) Survival rate of the mice 26 days post-injection of the AAVs. (C) Weight loss curve of the mice over the first five days post-surgery and the (D) individual weight loss on the fifth day post-surgery. The dotted line at 80% represents the threshold for euthanasia. (E) Latency to fall measured on an accelerating rotarod over three days with five trials per day. (F) Representative body motion traces measured over fifteen minutes of free moving mice in an openfield. (G) Total distance travelled of the mice measured in five frames of three minutes. (H) Representative body velocity and (I) acceleration curves measured in a period of two minutes. The shaded gray box represents locomotor bouts. (J) Quantification of the frequency of locomotor bouts. (K) Heatmaps of 195 randomly selected bouts one second prior and three seconds following the bouts initiation. The violet to yellow color palette represents the velocity from 0 to 200 mm/s. Scale bar = 0.5s. (L) Average velocity and (O) acceleration curve for a locomotor bout for each group. (M) Cumulative probability of locomotor bouts peak velocity, (N) duration, (P) peak acceleration, and (Q) peak deceleration. (R) Representative body motion traces measured over fifteen minutes in an elevated plus maze. (S) Quantification of the percentage of time spent in open arms in an elevated plus maze. (T) Quantification of the percentage of time spent grooming in a sucrose splash test. Filled dots represent the males and empty dots represent the females. Error bars represent the SEM. Pvalues were determined by a Student T-Test for S-T, one way ANOVA with Tukey’s post-hoc test for D, J, M-N, P-Q and two way ANOVA with Tukey’s post-hoc test for C, E, G. Pvalues annotated with an asterisk represents the comparison to No taCasp3 group, and hash represents the comparison to the Cre_ON_Flp_ON_-taCasp3 group. *P<0.05, **P<0.01, ***P<0.001, and ****P<0.0001.

We then assessed the role of CALB1^-^ and CALB1^+^ DA neurons on locomotor learning on an accelerating rotarod and recorded the latency to fall for untrained mice for 5 trials per day over three days. Mice with bilateral ablation of CALB1^+^ DA neurons showed no deficit in locomotor learning, displaying similar performance improvement than control mice (Figure3E). Contrastingly, bilateral or unilateral ablation of CALB1^-^ DA neurons had a severe impact on locomotor learning (Figure3E and FigureS3E). This poor performance following unilateral ablation could result from asynchrony, as demonstrated by the increase in ipsilateral turns and forepaw touch in the cylinder test (FigureS3F). Together, these results indicate that ablation of CALB1^-^, but not CALB1^+^ DA neurons, prevents locomotor learning.

To determine whether the effects of CALB1^-^ and CALB1^+^ DA neurons ablation extend beyond performance in a forced locomotor task, we next assessed how their removal affects voluntary movement in the openfield. Surprisingly, both CALB1^-^ and CALB1^+^ groups walked less in an openfield compared to control (Figure3F-G and FigureS3G-H). We then extracted locomotor bouts, defined as episodes in which the mouse’s body velocity exceeded 25mm/s for at least 600ms, thresholds chosen to exclude random movements from genuine locomotor events. We found that the frequency of locomotor bouts was significantly reduced when either CALB1^-^ and CALB1^+^ DA neurons were ablated (Figure3H-J and FigureS3I-K). Together, these results indicate that ablation of either CALB1^-^ or CALB1^+^ DA neurons impairs the initiation of voluntary movements.

We then looked at the vigor of ongoing movements by extracting the velocity and acceleration amplitudes within the locomotor bouts. Ablating bilaterally or unilaterally CALB1^-^ DA neurons resulted in a net decrease of the peak speed within the bouts (Figure3K-M and FigureS3L-N). However, we found that ablating bilaterally CALB1^+^ DA neurons resulted in a more pronounced deficit of the bout velocity amplitude than when the CALB1^-^ DA neurons were ablated (Figure3K-M). Minimal reductions in bouts length were observed when either population was ablated (Figure3N and FigureS3O). Finally, we found that both the bout’s peak acceleration and deceleration were significantly reduced when either CALB1^-^ or CALB1^+^ DA neurons were ablated (Figure3O-Q and FigureS3P-R). Since CALB1^+^ DA neurons are predominantly located in the VTA, a region known to modulate affective states including anxiety^71,72^, we considered the possibility that reduced locomotion might reflect increased anxiety rather than a motor deficit. To address this, we conducted an elevated plus maze test. No overall change was observed in time spent in the open arms, although a sex-specific analysis revealed a mild anxiety-like phenotype in females following ablation of CALB1^+^ DA neurons (P < 0.05) (Figure3R-S). Additionally, grooming behavior in the sucrose splash test was unaltered, suggesting that the observed motor deficits are unlikely to stem from an apathy-like phenotype (Figure3T). Together, these results indicate that the ablation of either CALB1^-^ or CALB1^+^ DA neurons impairs the initiation and vigor of movements.

### Chemogenetic inhibition of CALB1^-^ and CALB1^+^ DA neurons differently impairs the locomotor learning, initiation and vigor of ongoing movements

Ablation of DA neurons offers a simple way to assess the locomotor role of one subtype based on their presence or absence. However, the locomotor phenotype in CALB1^+^ DA neurons could result from the partial loss of CALB1^-^ DA neurons (Figure1D-G). Furthermore, following the ablation of a neuronal subtype, compensation can bias the behavioral outcome. To determine the role of these DA populations in intact circuits, we used chemogenetic to acutely inhibit either CALB1^+^ or CALB1^-^ DA neurons. In Calb1-ires-Cre/Dat-2A-Flp mice, we bilaterally injected either Cre_OFF_Flp_ON_ or Cre_ON_Flp_ON_ AAVs expressing the inhibitory receptor DREADD hM4Di along with a mCherry fluorescent reporter^73,74^. Control mice were injected with AAVs expressing an fDIO-mCherry reporter (Figure4A). Histologic analysis performed four weeks post-injection demonstrated that the infection spanned most of the VTA, SNc and RR (FigureS4A-B). Furthermore, quantification of mCherry reporter to TH^+^/CALB1^+^ and TH^+^/CALB1^-^ DA neurons confirmed the expression of the Cre_OFF_Flp_ON_ and Cre_ON_Flp_ON_ receptors to CALB1^-^ and CALB1^+^ DA neurons respectively (Figure4B and FigureS4C-D).

**Figure 4.**
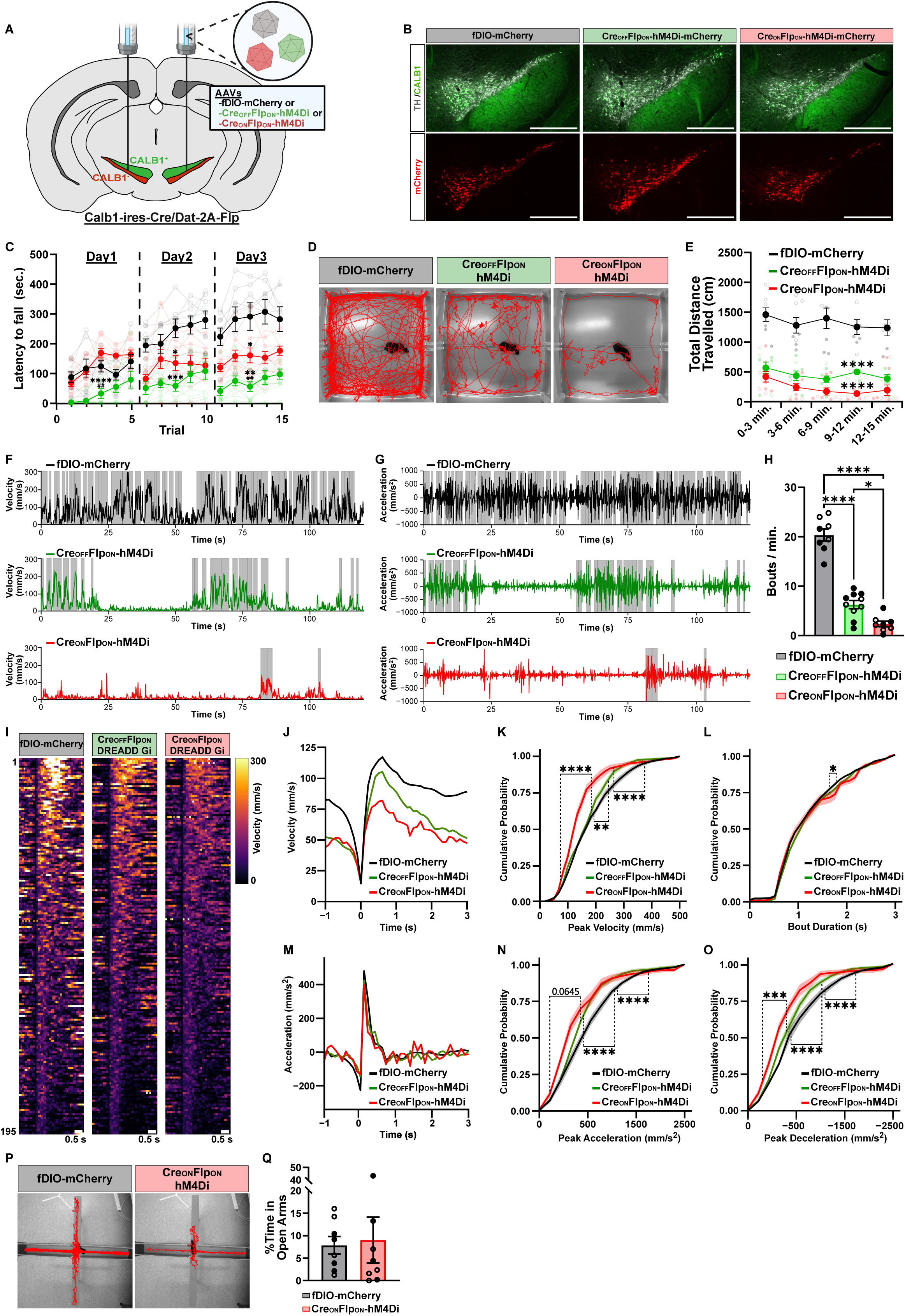
Bilateral acute inhibition of CALB1^-^ or CALB1^+^ DA neurons differently impairs locomotor learning, initiation, and vigor of ongoing movements. (A) Experimental strategy to bilaterally acutely inhibit either CALB1^-^ or CALB1^+^ DA neurons. Calb1-ires-Cre/Dat-2A-Flp mice were bilaterally injected of an AAV encoding either Cre_OFF_Flp_ON_ or Cre_ON_Flp_ON_-hM4Di inhibitory DREADD co-expressing the mCherry fluorescent reporter. (B) Representative images of midbrain sections co-stained for TH/CALB1 and mCherry fluorescent reporter. Scale bar = 500µm. (C) Latency to fall assessed on an accelerating rotarod over three days with five trials per day. (D) Representative body motion traces measured over fifteen minutes of free moving mice in an openfield. (E) Total distance travelled of the mice measured in five frames of three minutes. (F) Representative body velocity and (G) acceleration curves measured in a period of two minutes. The shaded gray box represents locomotor bouts. (H) Quantification of the frequency of locomotor bouts. (I) Heatmaps of 195 randomly selected bouts one second prior and three seconds following the bouts initiation. The violet to yellow color palette represents the velocity from 0 to 300 mm/s. Scale bar = 0.5s. Average velocity and (M) acceleration curve for a locomotor bout for each group. (K) Cumulative probability of locomotor bouts peak velocity, (L) duration, (N) peak acceleration, and (O) peak deceleration. (P) Representative body motion traces measured over fifteen minutes in an elevated plus maze. (Q) Quantification of the percentage of time spent in the open arms in an elevated plus maze. Filled dots represent the males and empty dots represent the females. Error bars represent the SEM. Pvalues were determined by a Student T-Test for S, one way ANOVA with Tukey’s post-hoc test for C-D, H, J, M-N, P-Q and two way ANOVA with Tukey’s post-hoc test for E, G. Pvalues annotated with an asterisk represents the comparison to fDIO-mCherry group, and hash represents the comparison to the Cre_ON_Flp_ON_-hM4Di group. *P<0.05, **P<0.01, ***P<0.001, and ****P<0.0001.

We next tested motor learning using the accelerating rotarod task following acute intraperitoneal injection of DCZ, a potent and selective DREADD hM4Di ligand^75,76^. Similar to the ablation, inhibition of CALB1^-^ DA neurons strongly impaired performance on all trial days (Figure 4C). In contrast, acute inhibition of CALB1^+^ DA neurons had no effect on performance on day one, but prevented performance improvement on days two and three (Figure 4C). These data indicate that while CALB1^-^ DA neurons are required for the initial acquisition of locomotor skills, CALB1^+^ DA neurons are essential for their retention. The inhibition of either CALB1^-^ or CALB1^+^ DA neurons impaired locomotor initiation (Figure4D-E), even though the frequency of locomotor bouts was further reduced by the inhibition of CALB1^+^ DA neurons (Figure4F-H). The vigor of ongoing movements was also impaired by the inhibition of either CALB1^-^ or CALB1^+^ DA neurons. Also, the bout peak velocity, acceleration, and deceleration were altered following the inhibition of both populations and were even further reduced for CALB1^+^ DA neurons (Figure4I-O). No change in the amount of time spent in the open arms was seen in an elevated plus maze following the acute inhibition of CALB1^+^ DA neurons (Figure4P-Q), suggesting that voluntary locomotion is unlikely biased by an anxiety-like phenotype. Together, these results demonstrate, for the first time, clear distinct roles of CALB1^+^ and CALB1^-^ DA neurons in locomotor learning, initiation and vigor of movements.

## DISCUSSION

Previous studies have established the molecular diversity of midbrain DA neurons, however, the functional role of these populations in locomotion has just begun to receive significant attention^25,28,29,33,35^. Here, we used chemogenetics and a novel intersectional taCasp3 vector to assess the role of CALB1 DA neurons, providing the first causal evidence that CALB1^+^ and CALB1^-^ DA neurons support distinct and complementary roles in locomotion. Our findings indicate that the bilateral ablation of CALB1^-^ DA neurons, but not CALB1^+^, is lethal, whereas both populations contribute to the initiation and vigor of movements. However, CALB1^-^ DA neurons are essential for motor performance, for instance on the accelerating rotarod, while CALB1^+^ DA neurons are required for the retention of acquired learning. Beyond our primary findings, we also observed a non-cell autonomous loss of CALB1^-^ DA neurons following CALB1^+^ ablation, coinciding with increased IBA1^+^ microglial density. While not the focus of our study, this raises the possibility that CALB1^-^ DA neurons may exhibit increased sensitivity to local inflammatory stress. Together, these findings demonstrate that CALB1 expression distinguishes populations of DA subtypes with complementary roles in locomotion.

Our findings have important implications for understanding the progression of motor symptoms in PD. While the resilience of CALB1^+^ DA neurons to degeneration in PD has been known for decades^21,33,53–58^, it remains surprising that no study has examined the behavioral consequence of their loss. Although the role of CALB1^-^ DA neurons in locomotion can be inferred from clinical observations, such assumptions are often confounded by synucleinopathy and degeneration of other brainstem nuclei. Our intersectional ablation approach offered a unique window for assessing the contribution of CALB1^-^ and CALB1^+^ DA neurons to locomotion in the absence of these confounds. We found that the ablation of CALB1^+^ DA neurons also impaired the initiation and vigor of movements. While this phenotype could be attributed to the observed loss of CALB1^-^ DA neurons, we report a similar locomotor deficit following acute inhibition of CALB1^+^ DA neurons, thereby confirming their direct contribution to locomotion. These findings contrast with previous reports showing no locomotor deficits following the ablation of VTA DA neurons^69,77–82^, regardless of the predominance of CALB1^+^ DA in this region^27,34^. Although the extent of the ablation and analysis pipeline differs from our investigation, taken together, it suggests that the CALB1^+^ DA neurons from the dorsal tier of the SNc, rather than those in the VTA, play a critical role in locomotion.

Our data offers a contrasting perspective on the effects of acute versus progressive DA loss, and implies that compensatory changes in PD and animal models should not be ignored. In particular, our work highlights how even partial perturbation of specific SNc subtypes leads to significant locomotor deficits. In PD, it is generally assumed that the onset of motor symptoms occurs when the degree of striatal DA loss exceeds 80%^83,84^. However, only a small minority of caudoputamen DA afference is from CALB1^+^ DA neurons^25,34^. Yet, their impact on locomotion is undeniable. Our results clearly demonstrated that the acute inhibition of a minority of SNc neurons can lead to a locomotor phenotype. Also, since the locomotor phenotype was more drastic following acute inhibition of CALB1^+^ neurons as compared to their ablation, it suggests that robust compensatory changes mitigate the observed phenotype following their permanent removal. Given that CALB1^+^ DA neurons encompass at least ten molecularly defined subtypes in mice^26,27^, it is possible that only a subset is responsible for the observed locomotor effects.

In addition, our findings provide important insights into the functional specialization of CALB1^+^ and CALB1^-^ DA neurons, likely driven by their unique connectivity. Past investigations suggested that the consolidation of motor learning is characterized by an engagement of the medium spiny neurons in the dorsomedial striatum, which is gradually relayed to the dorsolateral striatum as habits are formed^85–99^. Since the dorsomedial striatum receives input from CALB1^+^ DA neurons, it is consistent that acute inhibition of these neurons disrupted learning retention. Interestingly, we did not observe a similar effect following the ablation of CALB1^+^ DA neurons, suggesting that local compensatory mechanisms within the striatum may have masked this effect^100,101^. While the postsynaptic targets of CALB1^+^ and CALB1^-^ DA neurons can be reasonably inferred based on their projection patterns, far less is known about their specific presynaptic partners, despite previous efforts to broadly map mesocircuit connectivity^35,102,103^. If DA subtypes contribute differentially to motor learning, it is likely that they are integrated into parallel circuits that modulate action selection in subtype-specific ways. A more complete understanding of these networks will require future studies to delineate how each DA subtype is embedded within broader neural loops that support adaptive motor control.

We found that ablating or acutely inhibiting either CALB1^+^ or CALB1^-^ DA neurons impairs both peak acceleration and deceleration during locomotor bouts. While these findings are consistent with accumulating evidence that DA plays a permissive role in enabling vigorous movements^104–113^, it contrasts with the results of Azcorra et al.^25^, who reported that ANXA1^+^ and CALB1^+^ DA neurons activity correlates with acceleration and deceleration, respectively. Based on these prior findings, we anticipated that inhibition of CALB1^+^ neurons would preferentially impair deceleration. However, we observed that acceleration was similarly affected. Thus, our results add to previous findings highlighting a more general role of CALB1^+^ neurons in action selection that is not limited to deceleration. Thus, CALB1^+^ and CALB1^-^ DA neurons could differentially reinforce action selection during acceleration or deceleration^113,114^. Another possibility is that coordinated activity between these distinct DA subtypes is required for normal locomotion, and that acute perturbation of one population produces broader disruptions impacting both functions in a convergent manner. Future investigations could determine whether the distinct activity patterns of ANXA1^+^ and CALB1^+^ DA neurons are preserved following the inhibition of either population. Indeed, our results suggest that perturbing one subtype may exert profound effects on the activity of others. However, our study differs from the work of Azcorra et al. where recordings were limited to the dorsal striatum^25^, whereas our lesions/inhibition also affected DA release in the ventral striatum. Thus, it is possible that a specific DA neuron subtype, expressing CALB1 and innervating the caudoputamen, is causally tied to deceleration whose function was obscured by our broader perturbation.

The importance of phasic/dynamic DA release for locomotion has been recently questioned following reports that conditional knockout (cKO) mice for SYT1 or RIM1/2 in DA neurons do not exhibit locomotor deficits^115,116^. This contrasts with the marked impairment in openfield exploration we observed following chemogenetic inhibition of either CALB1^+^ or CALB1^-^ DA neuron populations. How can we explain that acute inhibition of DA neurons causes striking locomotor deficits, whereas deletion of SYT1 or RIM1/2, two proteins critical for DA release^117–121^, does not? Indeed, hM4Di is thought to affect behavior through a reduction of neuronal firing rates^73,74^, whereas both SYT1 and RIM1/2 are critical for pairing action potential with DA release, and their deletion virtually abolishes evoked DA and DA transients, respectively. One possibility is that the loss of RIM1/2 or SYT1 during development leads to compensatory mechanisms attenuating motor deficits in cKO animals^122,123^. This explanation, however, does not fully account for the fact that extracellular DA levels in the striatum are not altered in SYT1 cKO mice, despite these mice having a drastic reduction in evoked DA release. A second possibility is that chemogenetic inhibition of DA neurons impacts extracellular DA levels in the striatum through a SYT1-independent unidentified mechanism, an intriguing proposition that would need to be demonstrated experimentally.

Overall, our findings indicate that molecularly distinct DA neuronal subtypes encode different roles in locomotion and highlight the role of CALB1^+^ DA neurons, a contribution too long overlooked. Pending studies will be required to delineate further the distinct contribution of DA subtypes, their presynaptic regulation, and whether one DA population can compensate for the loss of another. These insights may ultimately inform novel circuit-based therapies aimed at reactivating or preserving residual DA neuron subtypes, offering promising avenues for mitigating motor symptoms of PD.

## RESOURCE AVAILABILITY

### Lead Contact

Jean-François Poulin

### Materials Availability

The INTRSECT taCasp3 will be deposited on Addgene.

### Data and Code Availability

The code generated for the extraction of locomotor bouts and analysis of elevated plus maze will be deposited on GitHub. All data reported in this paper will be shared by the lead contact upon request.

## ACKNOWLEDGEMENTS

The authors would like to thank Lucia Guerra for experimental support as well as George Sung, Kelvin Tiang, and Gabriella Talbot for feedback. CB received scholarships from Parkinson Canada, Fonds de recherche du Québec – Santé (FRQS), and Canadian Institutes of Health Research (CIHR), HB from HBHL and CIHR. JFP received funding from Parkinson Canada and CIHR (PJT-183760).

## AUTHOR CONTRIBUTIONS

CB designed, executed, analyzed experiments, and wrote the manuscript. CO, SD, HB, GT, ML, RM, MS, CC, CS, and SM executed, analyzed experiments, and provided feedback. RA developed the Dat-2A-Flp mice and edited the manuscript. JFP designed experiments and wrote the manuscript.

## SUPPLEMENTAL INFORMATION

Supplemental information will be deposited online.

---

**Figure S1.**
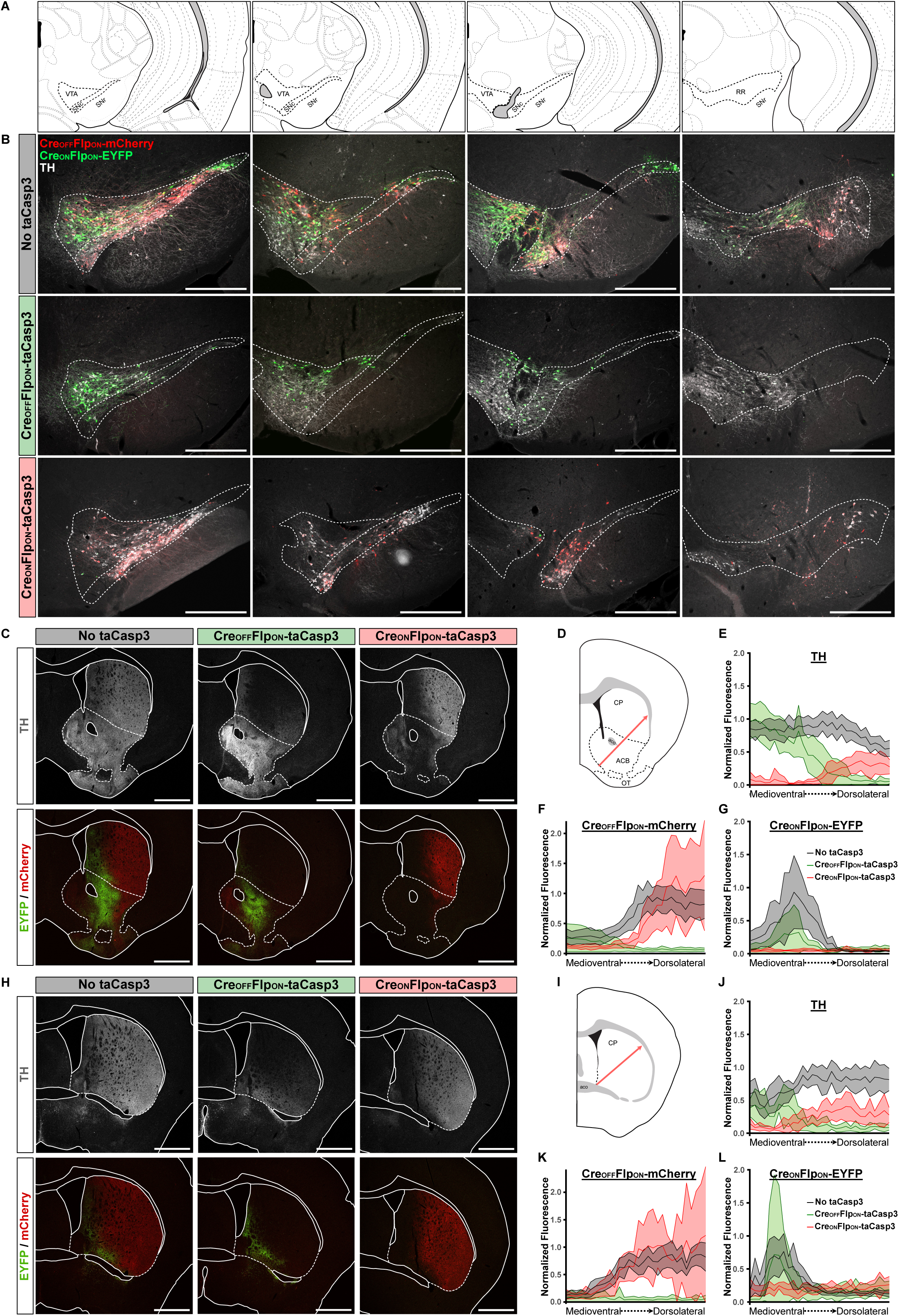
Extent of DA neurons ablation in the midbrain and striatal denervation. (A) Spatial location of the representative midbrain sections shown in B. The bold dotted lines represent anatomical regions where DA neurons were found. (B) Representative images of four rostro-caudal midbrain sections co-stained for EYFP/mCherry/TH. Scale bar = 500µm. (C) Representative images of rostral striatal sections stained for TH and EYFP/mCherry. Scale bar = 1000µm. (D) Cartoon showing the position of the arrow to quantify the level of denervation from the medioventral to dorsolateral segment of the rostral striatum. (E) Quantification in the rostral striatum of the medioventral to dorsolateral fluorescent level of TH, (F) Cre_OFF_Flp_ON_-mCherry, and (G) Cre_ON_Flp_ON_-EYFP relative to the control average maximal fluorescent intensity value. (H) Representative images of caudal striatal sections stained for TH and EYFP/mCherry. Scale bar = 1000µm. (I) Cartoon showing the position of the arrow to quantify the level of denervation from the medioventral to dorsolateral segment of the caudal striatum. (J) Quantification in the caudal striatum of the medioventral to dorsolateral fluorescent level of TH, (K) Cre_OFF_Flp_ON_-mCherry, and (L) Cre_ON_Flp_ON_-EYFP relative to the control average maximal fluorescent intensity value. Error bars represent the 95% Confidence Interval for E-G and J-L. Abbreviations: ACB = Nucleus accumbens, aco = anterior commissure, BST = Bed nuclei of the stria terminalis, CP = Caudoputamen, OT = Olfactory tubercle, RR = Retrorubral area, SNc = Substantia nigra pars compacta, SNr = Substantia nigra pars reticulata, VTA = Ventral tegmental area.

**Figure S2.**
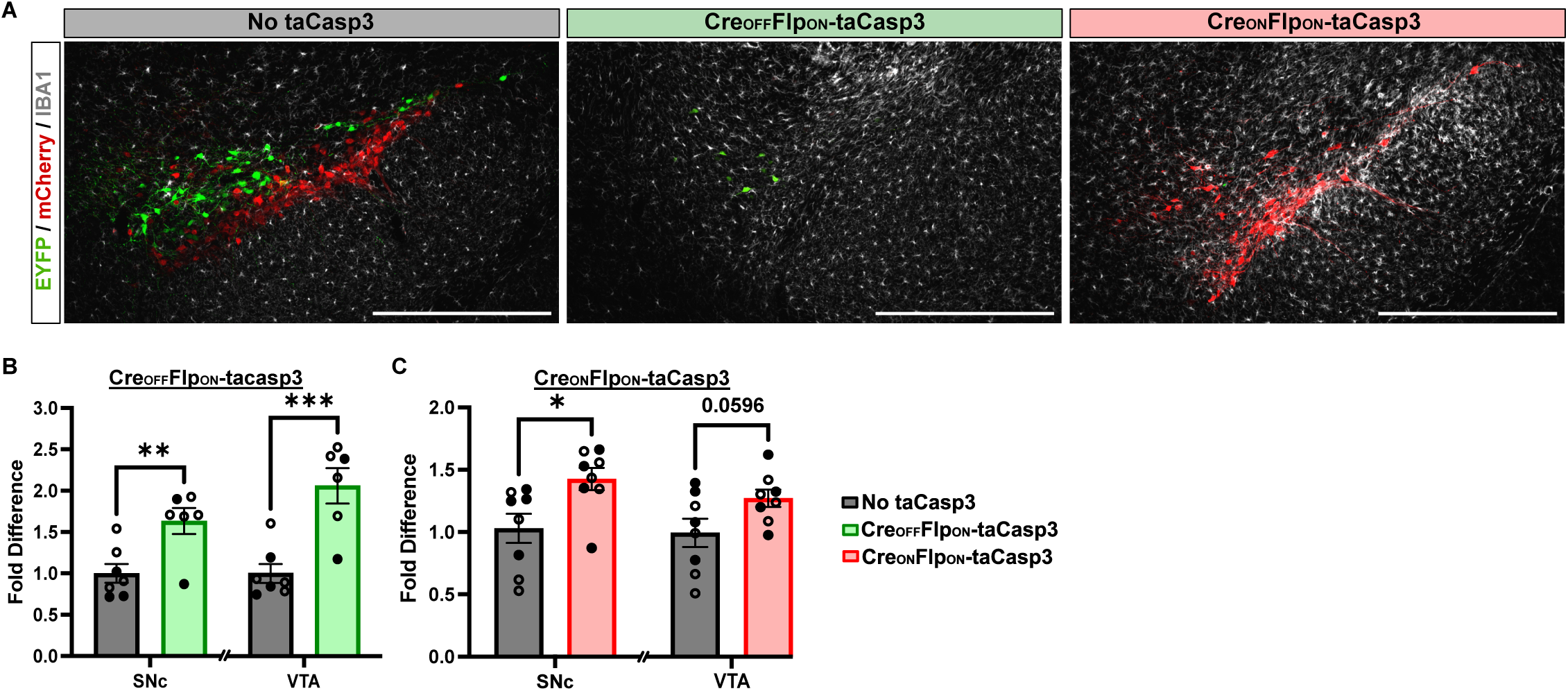
Increased density of IBA1^+^ microglia upon ablation of CALB1^-^ or CALB1^+^ DA neurons. (A) Representative images of midbrain sections co-stained for EYFP, mCherry and IBA1. Scale bar = 500µm. (B) Quantification of the amount of IBA1^+^ microglia in the SNc and VTA of Calb1-ires-Cre/Dat-2A-Flp mice injected with either Cre_OFF_Flp_ON_-taCasp3 or (C) Cre_ON_Flp_ON_-taCasp3 encoding AAVs. Filled dots represent the males and empty dots represent the females. Error bars represent the SEM. Pvalues were determined by a Student T-Test for B-C. *P<0.05, **P<0.01, ***P<0.001, and ****P<0.0001.

**Figure S3.**
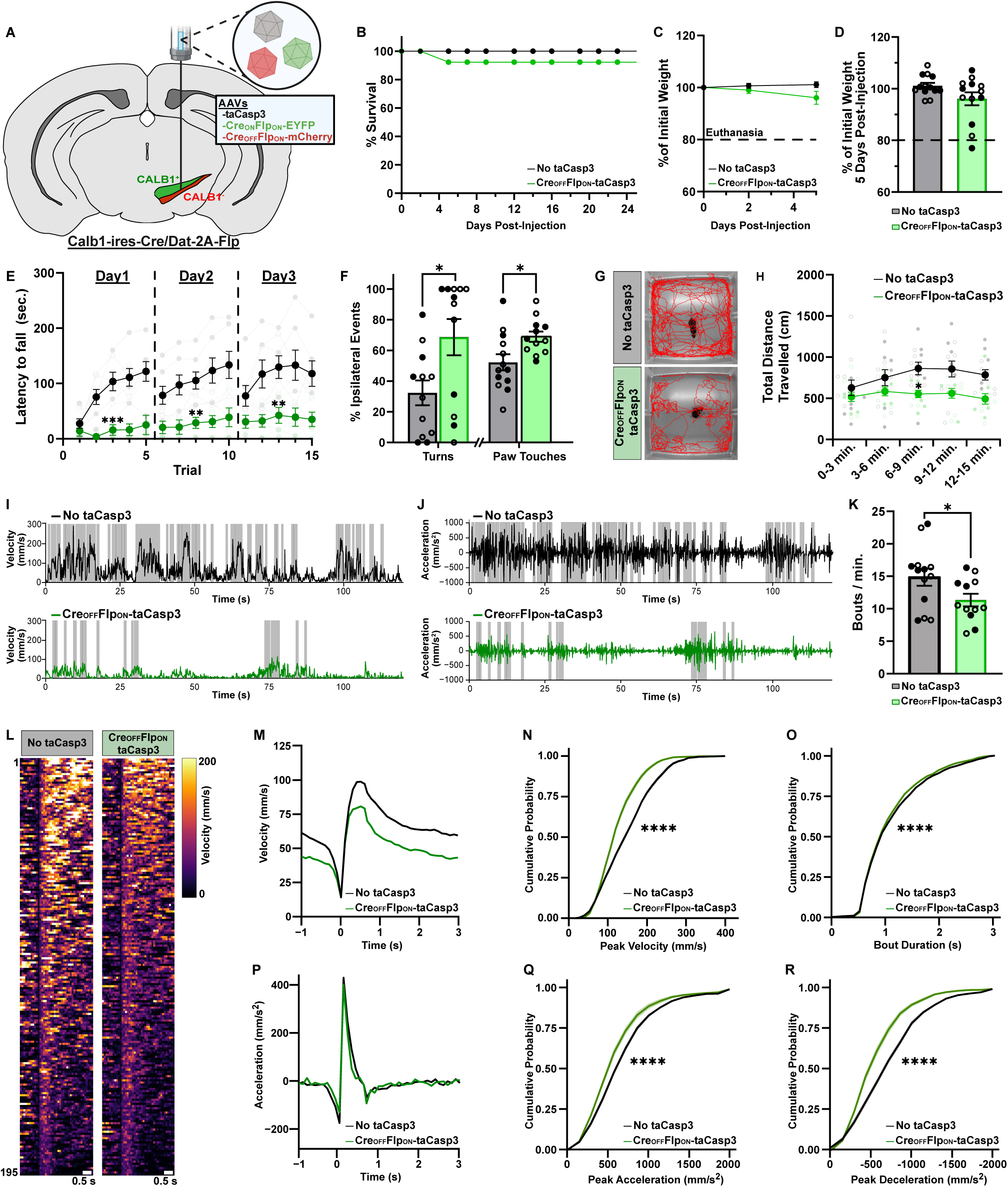
Unilateral ablation of CALB1^-^ DA neurons impairs locomotor learning, initiation, and vigor of ongoing movements. (A) Experimental strategy to unilaterally ablate CALB1^-^ DA neurons. Calb1-ires-Cre/Dat-2A-Flp mice were unilaterally injected with an AAV encoding Cre_OFF_Flp_ON_-taCasp3. Other AAVs encoding Cre_ON_Flp_ON_-EYFP and a Cre_OFF_Flp_ON_-mCherry were co-injected. (B) Survival rate of the mice 22 days post-injection of the AAVs. (C) Weight loss curve of the mice over the first five days post-surgery and the (D) individual weight loss on the fifth day post-surgery. The dotted line at 80% represents the threshold for euthanasia. (E) Latency to fall measured on an accelerating rotarod over three days with five trials per day. (F) Rate of ipsilateral turns and of use of the ipsilateral hindlimb measured in a cylinder test. (G) Representative body motion traces measured over fifteen minutes of free moving mice in an openfield. (H) Total distance travelled of the mice measured in five frames of three minutes. (I) Representative body velocity and (J) acceleration curves measured in a period of two minutes. The shaded gray box represents locomotor bouts. (J) Quantification of the frequency of locomotor bouts. (L) Heatmaps of 195 randomly selected bouts one second prior and three seconds following the bouts initiation. The violet to yellow color palette represents the velocity from 0 to 200 mm/s. Scale bar = 0.5s. (M) Average velocity and (P) acceleration curve for a locomotor bout for each group. (N) Cumulative probability of locomotor bouts peak velocity, (O) duration, (Q) peak acceleration, and (R) peak deceleration. Filled dots represent the males and empty dots represent the females. Error bars represent the SEM. Pvalues were determined by a Student T-Test for D, F, and K, one way ANOVA with Tukey’s post-hoc test for N-O, Q-R and two way ANOVA with Tukey’s post-hoc test for C, E, H. *P<0.05, **P<0.01, ***P<0.001, and ****P<0.0001.

**Figure S4.**
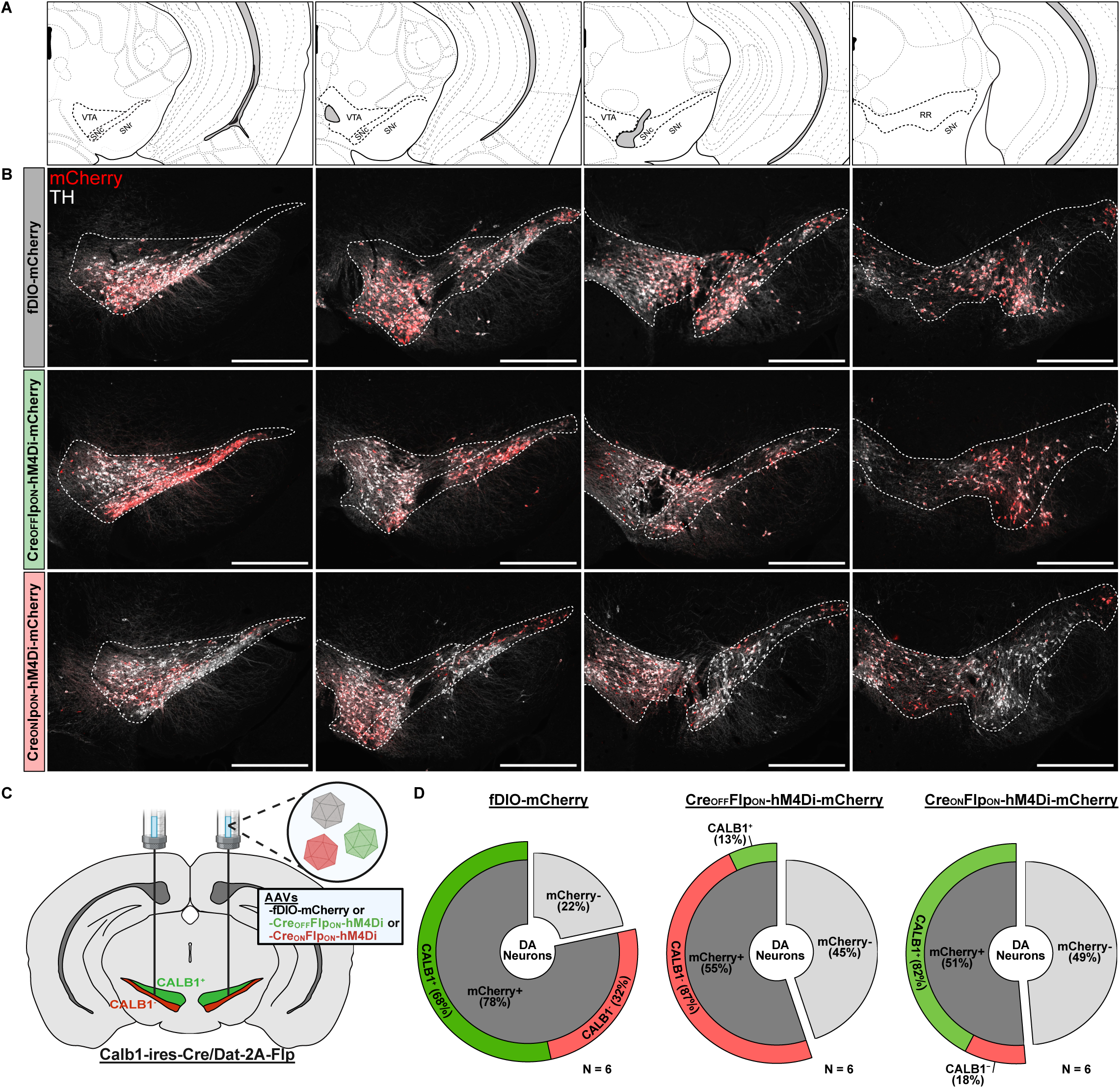
Infection coverage of inhibitory DREADD hM4Di in the midbrain. (A) Spatial location of the representative midbrain sections shown in B. The bold dotted lines represent anatomical regions where DA neurons were found. (B) Representative images of four rostro-caudal midbrain sections co-stained for mCherry/TH. Scale bar = 500µm. (C) Schematic of experimental strategy to bilaterally inhibit acutely either CALB1^-^ or CALB1^+^ DA neurons. Calb1-ires-Cre/Dat-2A-Flp mice were bilaterally injected of an AAV encoding either Cre_OFF_Flp_ON_ or Cre_ON_Flp_ON_ inhibitory DREADD_Gi_ chemogenetic receptor co-expressing the mCherry fluorescent reporter. (D) Donut charts of the proportion of mCherry^+^ and mCherry^-^ DA neurons. The proportion of mCherry^+^ DA neurons that are CALB1^+^ or CALB1^-^ are shown. Quantification was done in six individuals per group. Abbreviations: RR = Retrorubral area, SNc = Substantia nigra pars compacta, SNr = Substantia nigra pars reticulata, VTA = Ventral tegmental area.

## METHODS

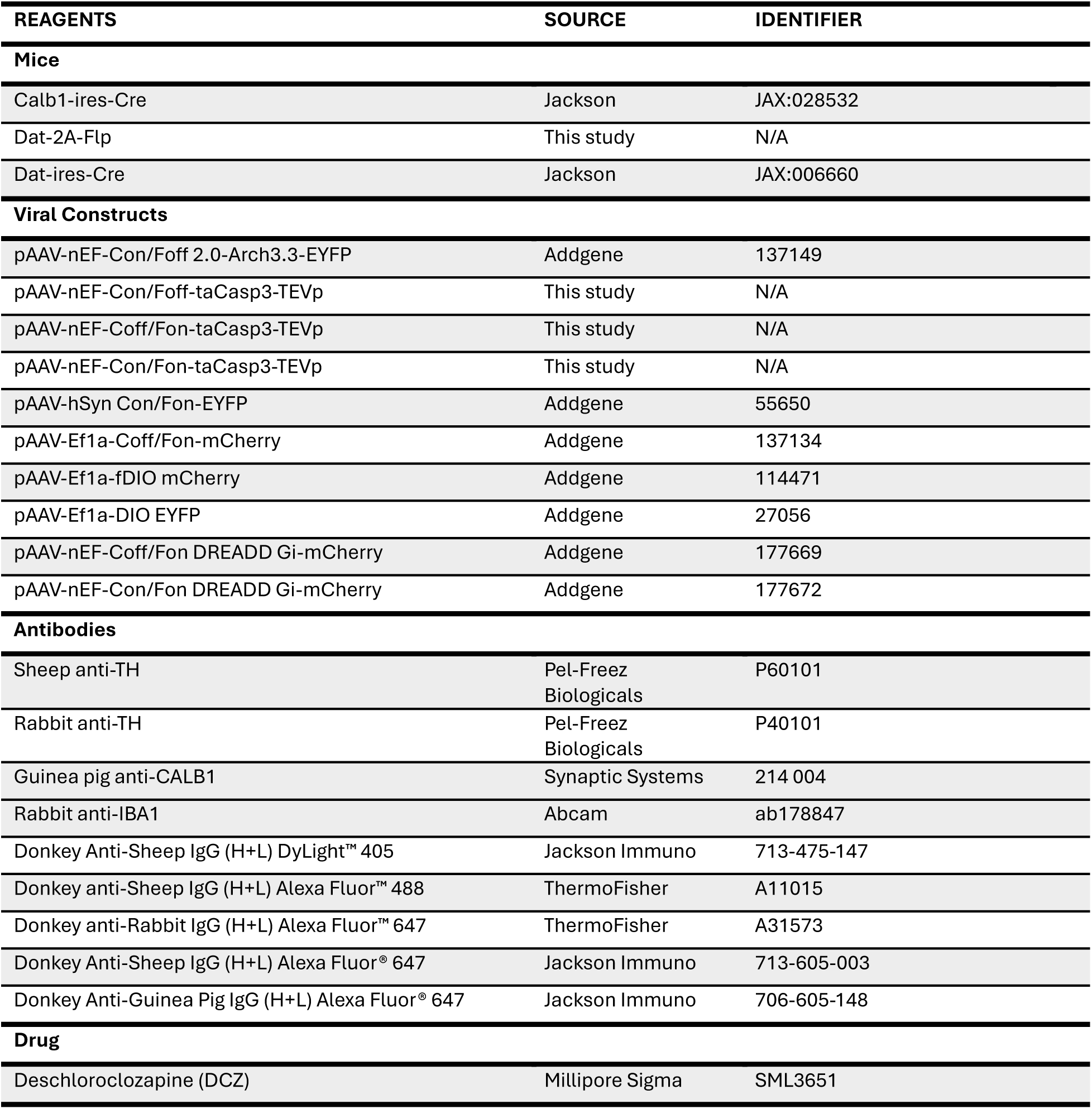
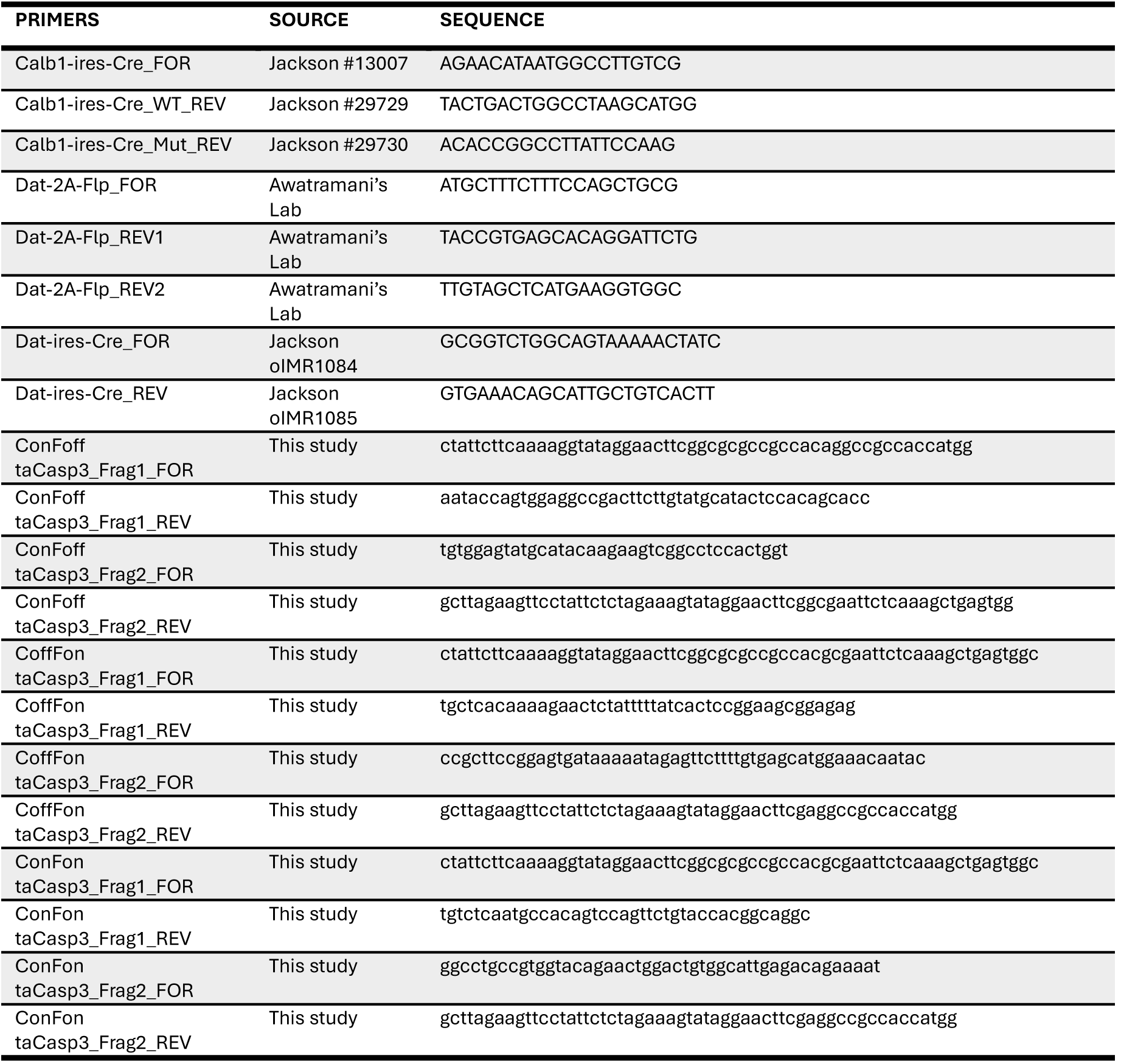

### Animals

All animals used in this study were maintained and cared following protocols approved by the Neuro Animal Care Committee from McGill University. Mice were housed at 20-22°C under 12 hours light/dark cycles with unlimited food and water source. Calb1-ires-Cre mice (Jackson, strain #028532) were maintained homozygous and crossed with heterozygous Dat-2A-Flp mice (generated at Northwestern University Transgenic and Targeted Mutagenesis Laboratory) for generation of experimental heterozygous Calb1-ires/Dat-2A-Flp mice. Heterozygous Dat-2A-Flp and Dat-ires-Cre mice (Jackson, JAX:006660) were also used for experiments. Both males and females were used in equal number for all experiments.

### Genotyping

Fourteen days post-natal mice tail samples were collected and genomic DNA was extracted with 200µL of 50mM NaOH and heated for at least 45 minutes at 95°C, neutralized with 50µL 1M of Tris (pH=8.0), and centrifuged for 6 minutes at 13,000 rpm. The DNA extracts were then processed by PCR using GoTaq® DNA Polymerase reaction kit (Promega, M3001). Calb1-ires-Cre mice were genotyped with the primers Calb1-ires-Cre_FOR, Calb1-ires-Cre_WT_REV, and Calb1-ires-Cre_Mut_REV. Dat-2A-Flp mice were genotyped with the primers Dat-2A-Flp_FOR, Dat-2A-Flp_REV1, and Dat-2A-Flp_REV2. Dat-ires-Cre mice were genotyped with the primers Dat-ires-Cre_FOR and Dat-ires-Cre_REV. All reaction cycles were carried at an initial denaturation of 5 minutes at 95°C, followed by 35 cycles of 30 seconds at 95°C with 30 seconds annealing at 60°C and 30 seconds at 72°C, then a final extension of 2 minutes at 72°C. The reactions were then run on agarose gel electrophoresis for identification of the WT and mutant mice.

### Cloning

The INTRSECT taCasp3 constructs were cloned similarly as described by Fenno et al. ^63,64^. Briefly, we designed synthetic oligonucleotides from Twist Bioscience, inserting introns CMV Towne Variant B(GenBank: M60321) and mouse IgE intron 3 (GenBank: X01857.1) into the coding sequence of pAAV-flex-taCasp3-TEVp (Addgene, Plasmid #45580)^65^. The target sites were chosen on the following criterias: first the two bases 5’ and two bases 3’ of the insertion match the exonic 5’ and 3’ consensus splice-site sequences, and second the insertion of the intron disrupts the reading frame disrupting the translation of a functional protein if the introns were not spliced. The likelihood that the inserted introns are functional was tested using NetGene2^124^. Controllers for Cre were incorporated by adding incompatible sites lox2722 and loxN^125^ to the introns, separated by a short spacer sequence^126^. Controllers for Flp were incorporated in the 5’ and 3’ ends of the taCasp3 coding sequence by adding the incompatible sites F3 and F5^127^ separated by the same spacer. The INTRSECT taCasp3 sequence was then synthesized as two complementary oligonucleotides using Twist Bioscience, amplified by PCR using 40 basepair overlapping primers. The reaction cycle consisted of an initial denaturation at 95°C for 4 minutes, then 30 cycles of denaturation at 94°C for 30 seconds, annealing at 55°C for 30 seconds, extension at 72°C for 45 seconds, and final extension at 72°C for 10 minutes. The amplicons were isolated on agarose gel using Monarch® Spin DNA Gel Extraction Kit (NEB, T1120). The plasmid pAAV-nEF-Con/Foff 2.0-Arch3.3-EYFP (Addgene, Plasmid#137149) was used as backbone by digestion with NcoI-HF (NEB, R3193) and NheI-HF (NEB, R0131), and the taCasp3 inserts were inserted by Gibson assembly^128^ using NEBuilder® HiFi DNA Assembly Cloning Kit (NEB, E5520). After transformation into NEB® 5-alpha Competent E. coli (NEB, C2987), miniprep was performed using QIAprep Spin Miniprep Kit (QIAGEN, 27104). Whole plasmid sequencing was performed by Plasmidsaurus using Oxford Nanopore Technology to confirm the achievement of the expected plasmidic sequence.

### Stereotaxic injection

Eight to twenty-five weeks old Calb1-ires-Cre/Dat-2A-Flp, Dat-ires-Cre or Dat-2A-Flp mice were used for surgeries. The mice were deeply anesthetized under isofluorane induction and subcutaneously injected with 20mg/kg carprofen and locally anesthetized with 0.1mL subcutaneous injection of a mix of 5mg/mL lidocaine and 2.5mg/mL bupivacaine. The mice were then headfixed and injected with 1000nL of virus at a rate of 200nL/min at the following coordinates: 3.15mm caudal, 1.2mm lateral, and 4.4 ventral to bregma. Following recovery, the mice were injected for three days post-surgery with 20mg/kg carprofen.

Calb1-ires-Cre/Dat-2A-Flp mice used for the ablation of CALB1^+^ or CALB1^-^ DA neurons were co-injected of 1×10^12^ genomic copies (GC)/mL of AAV2/DJ-Ef1a-Cre_OFF_Flp_ON_-mCherry, 1×10^12^ GC/mL of AAV2/DJ-hSYN-Cre_ON_Flp_ON_-EYFP, and 5×10^11^ GC/mL of either AAV2/9-nEF-Cre_OFF_Flp_ON_-taCasp3-TEVp or AAV2/9-nEF-Cre_ON_Flp_ON_-taCasp3-TEVp. Dat-2A-Flp mice used to test the specificity of the taCasp3 constructs in Flp expressing cells were co-injected with 1×10^12^ GC/mL of AAV2/9-EF1a-fDIO-mCherry, and 5×10^11^ GC/mL of either AAV2/9-nEF-Cre_OFF_Flp_ON_-taCasp3-TEVp or AAV2/9-nEF-Cre_ON_Flp_ON_-taCasp3-TEVp. Dat-ires-Cre mice used to test the specificity of the taCasp3 constructs in Cre expressing cells were co-injected with 1×10^12^ GC/mL of AAV2/9-EF1a-DIO-EYFP, and 5×10^11^ GC/mL of either AAV2/9-nEF-Cre_OFF_Flp_ON_-taCasp3-TEVp or AAV2/9-nEF-Cre_ON_Flp_ON_-taCasp3-TEVp. Calb1-ires-Cre/Dat-2A-Flp mice used for the acute silencing of CALB1^+^ or CALB1^-^ DA neurons were co-injected of either 1×10^12^ GC/mL of AAV2/9-EF1a-fDIO-mCherry, AAV2/9-nEF-Cre_OFF_Flp_ON_-DREADD_Gi_-mCherry or AAV2/9-nEF-Cre_ON_Flp_ON_-DREADD_Gi_-mCherry. All AAVs were produced by the Molecular Tools Platform from Laval University, and eluted in PBS with 320mM NaCl, 5% (w/v) D-Sorbitol, and 0.001% (v/v) F68.

### Rotarod

Rotarod was used to assess the locomotor learning and performance of the mice. All mice were assessed three weeks post-injection of the AAVs. Prior to the trials, the mice were habituated to the room for at least 30 minutes. The latency to fall of the mice was assessed over three days with five trials per day. For CALB1^+^ or CALB1^-^ DA neurons ablated mice, the latency to fall on an accelerating wheel (Harvard Apparatus, 76-0770) passing from 4 to 40 rotations per minute in five minutes was measured. For mice with chemogenetic inhibition of CALB1^+^ or CALB1^-^ DA neurons, trials were also performed three weeks post-injection of the AAVs. Each day, the mice received an intraperitoneal injection 0.1mL per 20g of body weight of a solution of 0.02mg/mL deschloroclozapine (DCZ) at least 30 minutes prior to the first trial, and latency to fall was measured on an accelerating wheel passing from 4 to 40 rotations per minute in ten minutes.

### Body motion tracking in an openfield

Mice voluntary movements were evaluated using the openfield test to assess the impact of neuronal ablation or chemogenetic inhibition on movements initiation and vigor. Behavioral assessments were performed three weeks following AAV injection. Prior to testing, mice were habituated to the room for at least 30 minutes. Mice with chemogenetic inhibition of either CALB1^+^ or CALB1^-^ DA neurons received an intraperitoneal injection 0.1mL per 20g of body weight of a solution of 0.02mg/mL DCZ at least 30 minutes before the trial. The mice were placed in a 15.5" × 16" openfield arena and recorded for 15 minutes using an ELP 1080P Day/Night Vision USB Camera (ShenZhen Ailipu Technology Co., ELP-USBFHD05MT-KL36IR).

Body motion was tracked using DeepLabCut, version v2.3^129^. A custom-trained neural network was used to detect key anatomical landmarks, including the nose, head, ears, base and mid-point of the tail, body humps, and hips. To reduce tracking variability during periods of immobility, only the body hump point was used for motion analysis.

Tracking data were processed using custom R scripts with the packages ggplot2, tidyverse, imager, and viridis. Frames were filtered using a confidence threshold of 0.9, with low-confidence detections excluded. The videos were downsampled to 1 frame every 100ms.

Total distance traveled was calculated by summing the Euclidean distances between consecutive body hump positions in 3-minute intervals. Velocity was estimated as the Euclidean distance between two consecutive frames divided by the time elapsed. Acceleration was calculated as the change in velocity between consecutive frames divided by the time interval.

To exclude random movements from genuine locomotor events, locomotor bouts were defined as periods during which the velocity of the body hump exceeded 25 mm/s for at least 600 ms. The frequency of locomotor bouts was computed by dividing the number of bouts by the total time of the video. The distributions of peak velocity, acceleration, and deceleration during bouts were analyzed by binning their amplitudes into 31 bins for velocity and 16 bins for acceleration/deceleration, and expressing each as a cumulative probability relative to the total number of bouts.

### Cylinder test

Mice with unilateral ablation of CALB1^-^ DA neurons were placed in a transparent cylinder to evaluate ipsilateral bias in hindlimb use during wall contacts, as well as ipsilateral body turns. Prior to testing, mice were habituated to the testing room for at least 30 minutes. Each mouse was recorded for 10 minutes from below using an ELP 1080P Day/Night Vision USB Camera (ShenZhen Ailipu Technology Co., ELP-USBFHD05MT-KL36IR). Behavioral analysis was performed manually using BORIS^130^.

### Elevated plus maze

Mice with either bilateral ablation or chemogenetic inhibition of CALB1^+^ DA neurons underwent an elevated plus maze to assess the presence of an anxiety-like phenotype. Prior to testing, mice were habituated to the testing room for at least 30 minutes. Then, the mice were placed to the middle of the maze and recorded for 15 minutes using an ELP 1080P Day/Night Vision USB Camera (ShenZhen Ailipu Technology Co., ELP-USBFHD05MT-KL36IR). The body motion of the mice were tracked using DeepLabCut as described above. The percentage of time that the complete body of the mice spent in the open arms was extracted.

### Sucrose splash test

Sucrose splash test was performed on the mice with bilateral ablation of CALB1^+^ DA neurons to assess the presence of apathy-like phenotype. Prior to testing, mice were habituated to the testing room for at least 30 minutes. Then, the mice were placed into a clean transport cage, and sprayed once with a 10% (w/v) sucrose solution, and recorded for 5 minutes using an ELP 1080P Day/Night Vision USB Camera (ShenZhen Ailipu Technology Co., ELP-USBFHD05MT-KL36IR). The amount of time the mice spent grooming was analyzed manually using BORIS.

### Perfusion

Four weeks post-injection of the AAVs, the mice were perfused for brain collection. The mice were deeply anesthetized and maintained under isofluorane induction, and intracardiacally perfused at a rate of 6mL/min with 40mL of room temperature 1X PBS followed by 40mL of ice-cold 4% (w/v) PFA in 1X PBS. The brains were then extracted and fixed overnight at 4°C, and immerged in 30% (w/v) sucrose in 1X PBS until completely sunk. Then, the brains were frozen in OCT and preserved to −80°C until processed for histology.

### Histology

The brains were sliced coronally in 25µm thick sections using a Leica CM1850 Cryostat, and preserved in a cryoprotectant solution containing 11.5mM NaH_2_PO_4_, 38.5mM Na_2_HPO_4_, 30% (v/v) RNAse free ethylene glycol and 20% (v/v) RNAse free glycerol. For quantification of the extent of ablation or to assess the infection rate of hM4Di, six midbrain sections at about 200µm intervals were picked to cover slices 79 to 91 of the Allen Institute Coronal Atlas. For striatal sections, four sections corresponding to the slices 44, 49, 53, and 58 of the Allen Institute Coronal Atlas were picked.

The sections were picked, washed three times for 10 minutes in 1X PBS, and blocked for 30 minutes at room temperature with shaking in 4% (v/v) normal donkey serum (NDS) + 0.5% (v/v) Triton-X100 in 1X PBS blocking buffer. Then, the sections were incubated overnight at 4° C with shaking in blocking buffer with the following primary: sheep anti-TH (Pel-Freez Biologicals, p60101, 1:500-1000), rabbit anti-TH (Pel-Freez Biologicals, P40101, 1:1000), guinea pig anti-CALB1 (Synaptic Systems, 214 004, 1:1000), and rabbit anti-IBA1 (Abcam, ab178847, 1:200). The sections were then washed three times 10 minutes in 1X PBS + 0.05% (v/v) Tween20, and incubated for two hours at room temperature with shaking in blocking buffer with the following secondary antibodies: Donkey Anti-Sheep IgG (H+L) DyLight™ 405 (Jackson Immuno, 713-475-147, 1:200), Donkey anti-Sheep IgG (H+L) Alexa Fluor™ 488 (ThermoFisher, A11015, 1:250), Donkey anti-Rabbit IgG (H+L) Alexa Fluor™ 647 (ThermoFisher, A31573, 1:1250), Donkey Anti-Sheep IgG (H+L) Alexa Fluor® 647 (Jackson Immuno, 713-605-003, 1:200), and Donkey Anti-Guinea Pig IgG (H+L) Alexa Fluor® 647 (Jackson Immuno, 706-605-148, 1:200). Then, the sections were washed three times 10x minutes in 1X PBS, and mounted in Anti-Fade Fluorescence Mounting Medium (Abcam, ab104135).

Sections were imaged using a Nikon Eclipse Ti2 inverted confocal microscope. All images from the same immunohistochemistry batch were acquired using identical imaging parameters. For cell body quantification, a segmented model was trained using the NIS-Elements NIS.ai software. The training was based on at least 5,000 iterations of manually labeled cells by a human experimenter from a binary layer. The trained model was then applied to the images, after which false positives and negatives were manually corrected. The number of binary objects was subsequently quantified within the designated regions of interest (ROI).

### Striatal denervation spectra analysis

Confocal images of striatal sections were analyzed using a custom ImageJ macro and R script. Briefly, images were opened in ImageJ, channels were split, and arrows were drawn along the dorsoventral, mediolateral, and medioventral–dorsolateral axes. Pixel intensity values along these axes were extracted and exported to a spreadsheet. Additionally, two ROIs were placed in the cortex and one in the septum, and the mean intensity values from these ROIs were saved in a separate spreadsheet.

To correct for background noise, pixel intensity values along the measured axes were individually adjusted by subtracting the mean signal from the cortical and septal ROIs. Any pixel value lower than the mean noise signal was set to zero. Corrected values were then averaged into 30 bins and normalized to the maximum intensity of the average curve from the control group, processed within the same immunohistochemistry batch. Normalized curves from individual animals were pooled based on their corresponding slice levels, and 95% confidence intervals were computed to assess differences in fluorescence intensity along the measured axes.

